# Consensify: a method for generating pseudohaploid genome sequences from palaeogenomic datasets with reduced error rates

**DOI:** 10.1101/498915

**Authors:** Axel Barlow, Stefanie Hartmann, Javier Gonzalez, Michael Hofreiter, Johanna L. A. Paijmans

**Author notes:** Corresponding authors: AB, JLAP.

## Abstract

A standard practise in palaeogenome analysis is the conversion of mapped short read data into pseudohaploid sequences, typically by selecting a single high quality nucleotide at random from the stack of mapped reads. This controls for biases due to differential sequencing coverage but it does not control for differential rates and types of sequencing error, which are frequently large and variable in datasets obtained from ancient samples. These errors have the potential to distort phylogenetic and population clustering analyses, and to mislead tests of admixture using D statistics. We introduce Consensify, a method for generating pseudohaploid sequences which controls for biases resulting from differential sequencing coverage while greatly reducing error rates. The error correction is derived directly from the data itself, without the requirement for additional genomic resources or simplifying assumptions such as contemporaneous sampling. For phylogenetic analysis, we find that Consensify is less affected by branch length artefacts than methods based on standard pseudohaploidisation, and it performs similarly for population clustering analysis based on genetic distances. For D statistics, Consensify is more resistant to false positives and appears to be less affected by biases resulting from different laboratory protocols than other available methods. Although Consensify is developed with palaeogenomic data in mind, it is applicable for any low to medium coverage short read datasets. We predict that Consenify will be a useful tool for future studies of palaeogenomes.

## 1. INTRODUCTION

The recovery of nuclear genomic data from ancient biological material – i.e. palaeogenomic data – is typically challenged by high levels of contamination, a low abundance of ancient nucleic acids, and the physical properties of the molecules themselves, such as short fragment length and the presence of miscoding and blocking lesions (Briggs et al., 2007; Brotherton et al., 2007; Heyn et al., 2010; Hofreiter, Jaenicke, Serre, Haeseler, & Pääbo, 2001). Therefore, it can be assumed that the per nucleotide expense of data recovery from ancient samples will be considerably greater than for an equivalent living organism. Disregarding financial costs, there may also be physical limits on data recovery as sufficient template molecules may simply not be present for high coverage palaeogenome sequencing of some ancient samples. As a result, published palaeogenome datasets have typically been low coverage (Barlow et al., 2018; Green et al., 2010, 2006; Orlando et al., 2013; Palkopoulou et al., 2018; Skoglund, Ersmark, Palkopoulou, & Dalén, 2015), while high coverage datasets are comparatively scarce (Meyer et al., 2012; Palkopoulou et al., 2015; Prüfer et al., 2017).

Low coverage datasets present a particular challenge for data analysis. Standard SNP calling approaches, especially involving the identification of heterozygous positions, are likely to be error-prone when applied to low coverage palaeogenome data, although methods have been developed for bypassing the problems to some extent (Kousathanas et al., 2017). Sophisticated methods for estimating SNPs also exist but these are only applicable for specific datasets such as human SNPs, e.g. (Schraiber, 2018). Despite these more complex approaches, arguably the most frequently used approach has been to sample a single high quality nucleotide from the read stack for each position of the reference genome (Green et al., 2010). Assuming equal rates of sequencing and mapping errors among samples, these so-called pseudohaploid sequences effectively downsample all datasets to 1x coverage; thus, normalising the rate of errors among datasets. However, differential sequencing and mapping errors among palaeogenomic datasets may exist and be large, due to variability in fragment length distributions, levels of cytosine deamination, and laboratory artefacts (Barlow et al., 2016).

Differential errors in pseudohaploid sequences have the potential to confound both phylogenetic and population clustering (e.g. PCA, principal coordinates analysis) analyses. Increased error rates in some datasets, for example ancient compared to modern datasets, are likely to manifest as an excess of singleton sites. For phylogenetic analysis, this will result in an increase in the lengths of terminal branches leading to the high-error individuals. Although the internal topology of the tree is less likely to be affected, it is feasible that for more complex analyses that involve constraining the tip ages, the affected lineages could be artefactually pushed to more basal positions in the tree. For population clustering analysis, it is feasible that errors could dominate the variability leading to individuals clustering by error rate rather than ancestry. Furthermore, both absolute and relative estimates of diversity or divergence are likely to be confounded if applied to datasets with substantial differences in error rates.

Several methods have been employed to reduce the effect of differential errors on phylogenetic and clustering analyses. A major cause of sequencing errors in palaeogenomic datasets is cytosine deamination, which manifests as C→T (and in some cases additionally G→A) substitutions (Briggs et al., 2007; Brotherton et al., 2007; Hofreiter et al., 2001). Although the standard practise of excluding transition sites for the purpose of analysis is an effective means of dealing with this source of errors, non-clocklike evolution observable in published phylogenetic trees (e.g. Barlow et al., 2018) suggests that transversion-based errors in palaeogenomic datasets are also appreciable. A potentially useful method is to remove all singletons from the dataset prior to analysis (e.g. Westbury et al., 2018). This can be effective if clades are reasonably and equally sampled. However, undersampled divergent lineages will experience a removal of private “real” substitutions and consequently exhibit terminal phylogenetic branch shortening artefacts following singleton removal, as well as potentially fail to form distinct populations in population clustering analyses.

Another class of analyses that may be confounded by differential errors in pseudohaploid sequences are tests of admixture, such as the frequently used D statistic (Durand, Patterson, Reich, & Slatkin, 2011; Green et al., 2010). In its original form, the D statistic uses standard pseudohaploid sequences from two closely related individuals (P1, P2), a third individual representing a candidate admixing lineage (P3), and a fourth individual (P4) that represents the outgroup. Their phylogeny is: (((P1, P2), P3), P4). For biallelic sites, alleles sampled in the outgroup are assumed to be ancestral (A) and the alternate allele is therefore derived (B). The D statistic is the difference in the frequencies of sites where P2 and P3 share a derived allele not found in P1 (so called ABBA sites) and those where P1 and P3 share a derived allele not found in P2 (so called BABA sites), normalised for the number of observations. D scales between −1 and +1 with positive values (excess of ABBA sites) suggesting admixture between P2 and P3 subsequent to the divergence of P1 and P2, and negative values (excess of BABA sites) suggesting admixture between P1 and P3, subsequent to the divergence of P1 and P2.

Although the D statistic provides a powerful test of admixture, it assumes that alleles are sampled without error (Durand et al., 2011). Differential error rates between P1 and P2 individuals present a particular problem, however. By example, if P2 is ancient and P1 is modern, increased errors in the ancient dataset will cause a proportion of BBBA sites to be converted to D statistic informative BABA sites. As a result, the high error P2 individual will appear increasingly unadmixed relative to P1 (Barlow et al., 2018; Orlando et al., 2013). This effect is magnified with more divergent outgroups, since they will possess more private alleles. Such effects have been observed by analysis of empirical datasets, where the D value can be shifted from significantly positive to significantly negative by using increasingly diverged outgroups (Barlow et al., 2018). Recently, efforts have been made to apply a statistical correction to these artefacts. The extended D statistic (Soraggi, Wiuf, & Albrechtsen, 2017), rather than standard pseudohaploidisation, makes use of the complete read stack and can further apply a correction to error rates estimated by comparison to data from a high quality “error free” individual. This method assumes that an excess of singletons in the test dataset relative to the error free individual is attributed to error, and can be use to correct the observed ABBA and BABA counts. In theory, this provides a true error correction by normalising error rates to that of the error free individual. An implicit assumption, however, is that all individuals are sampled contemporaneously. If the test dataset is from an individual that is appreciably older than the error free individual, then error rates may be underestimated as the ancient lineage has has less time for substitutions to accrue.

In this study, we present Consensify, a method for reducing error rates in pseudohaploid sequences generated from palaeogenomic and other low to medium coverage datasets. The error reduction is derived directly from the data itself, and does not require additional resources such as a high coverage data from a close relative, nor does it require simplifying assumptions such as a strict molecular clock or contemporaneous sampling. We show that Consensify brings qualitative improvement for phylogenetic and population clustering analyses. For admixture tests, we also demonstrate by simulation that Consensify is more resistant to false positives than other available methods, and is generally more conservative than other methods when applied to real-world empirical examples. Consensify thus represents a useful tool for future studies of palaeogenomes.

## 2. MATERIALS AND METHODS

### 2.1 The Consensify method

Consensify is a simple method for generating consensus pseudohaploid sequences from sequencing reads that are mapped to a reference genome. For each position, three nucleotides are extracted from the read stack at random. If two of the reads agree, then that base is retained. If only two reads are present, but they agree, then that base is also retained. If no two reads agree, then an N is entered for that position. If coverage is < 2, or above a maximum depth specified by the user, then an N is entered for that position. An example is shown below. The table summarises a read stack by the number of bases observed in columns (totA, totC, totG, totT) at each position of the reference genome (represented by sequential rows). The Consensify sequence for this read stack would be TGNAC.

| totA | totC | totG | totT |
| --- | --- | --- | --- |
| 0 | 0 | 1 | 2 |
| 0 | 0 | 2 | 0 |
| 1 | 0 | 0 | 1 |
| 4 | 0 | 0 | 0 |
| 0 | 4 | 1 | 0 |

To explore the statistical properties of the Consensify method, we considered a simple model of sequencing error assuming equal genomic base composition and assuming that sequencing errors occur with equal probability across all possible nucleotide combinations. This error model is conceptually identical to the JC69 model of nucleotide substitution (Jukes & Cantor, 1969). In this model, the global error rate can be summarised by a single variable (*P*errorGlobal). The probability of observing any base as an error (*P*errorBase) is therefore:

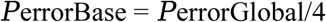

For any homozygous position, the probability of observing the correct base (*P*correctHom) in a single sequencing read is:

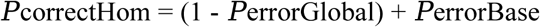

The last term in the equation reflects that, according to the model, it is possible for a sequencing error to replace a base with the identical base.

For heterozygous positions, the probability of observing a correct base is higher, since an error may convert a base to either allele, thus:

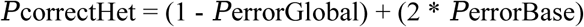

Sampling three nucleotides mapped to a single genomic position has 64 possible outcomes. By applying this model of sequencing error it is possible to calculate the probability of observing each outcome given a particular genotype. Summing the relevant probabilities allows the probability of observing a correct base, missing data (N), or an incorrect base, using the Consensify method (Supplementary Information). We calculated expected error rates for Consensify assuming this model and compared them with expectations for standard pseudohaploidisation.

### 2.2 Test datasets

We tested the Consensify method using published Illumina paired-end sequencing datasets of bears (Barlow et al., 2018; Benazzo et al., 2017; Cahill et al., 2013, 2015; Kumar et al., 2017). These comprised three brown bears *(Ursus arctos)*, two polar bears *(Ursus maritimus)*, an Asiatic black bear *(Ursus thibetanus)* and four Late Pleistocene cave bears *(Ursus spelaeus* complex). The relationship of these clades is (black,(cave,(polar,brown))). The cave bear datasets represent four taxa defined based on morphology and mitochondrial DNA, and their relationship is *(kudarensis,(eremus, (spelaeus,ingressus)))* (Barlow et al., 2018). Full details of the datasets analysed are provided in Table 1.

**Table 1.** Details of datasets included in this study.

| Dataset | Taxon | Data type | Reference | Mapped Gb<sup>1</sup> | Consensify sites<sup>2</sup> |
| --- | --- | --- | --- | --- | --- |
| E-VD-1838 | cave bear (*spelaeus*) | ancient single-stranded | Barlow et al. 2018 | 4.55215 | 971,153,181 |
| GS136_ds | cave bear (*ingressus*) | ancient double-stranded | Barlow et al. 2018 | 3.72732 | 519,642,820 |
| GS136_ss | cave bear (*ingressus*) | ancient single-stranded | this study | 6.94074 | 1,266,005,835 |
| WK01 | cave bear (*eremus*) | ancient single-stranded | Barlow et al. 2018 | 6.12884 | 1,210,933,302 |
| HV74 | cave bear (*kudarensis*) | ancient single-stranded | Barlow et al. 2018 | 3.76075 | 869,048,390 |
| 191Y | brown bear (Slovenia) | modern | Barlow et al. 2018 | 6.99668 | 1,088,167,390 |
| SRS779830 | brown bear (Sweden) | modern | Cahill et al. 2015 | 6.13821 | 1,076,186,512 |
| SRR5878360 | brown bear (Italy) | modern | Benazzo et al. 2017 | 17.51220 | 1,553,722,573 |
|  |  | simulated ancient (35 bp, damage) |  | 7.42132 | 1,293,235,675 |
|  |  | simulated ancient (35 bp, error) |  | 7.55782 | 1,297,732,079 |
|  |  | simulated ancient (35 bp, damage, error) |  | 7.36212 | 1,291,668,999 |
|  |  | simulated ancient (50 bp, damage, error) |  | 10.51792 | 1,109,680,441 |
| SRS412584 | polar bear | modern | Cahill et al. 2013 | 6.81197 | 1,118,483,213 |
| SRS412585 | polar bear | modern | Cahill et al. 2013 | 6.02194 | 1,025,241,632 |
| ERS781634 | Asiatic black bear | modern | Kumar et al. 2017 | 13.43641 | 1,523,577,868 |

<sup>1</sup>Gb data successfully mapped to panda reference genome assembly
<sup>2</sup>Number of sites successfully called using Consensify

The cave bear datasets are palaeogenomic datasets that feature the typical properties of ancient DNA (Barlow et al., 2018). The vast majority of sequences for three of the published cave bear datasets were generated from sequencing libraries prepared using a method based on single-stranded DNA (Gansauge & Meyer, 2013), whereas the fourth dataset *(ingressus*, GS136_ds) was generated from sequencing libraries prepared using a method based on double-stranded DNA (Meyer & Kircher, 2010). For this study, we additionally prepared a single-stranded library from DNA extracted from the same petrous bone of the *ingressus* cave bear previously sequenced from only double-stranded libraries, using the method outlined in (Gansauge & Meyer, 2013) exactly following the procedure described in (Basler et al., 2017) and sequenced it on an Illumina NextSeq 500 platform returning 75bp dual-indexed single-end reads, following the procedure described in (Paijmans et al., 2017). These datasets allow a direct comparison of the effect of the method of library preparation on downstream analyses.

Processing of sequence data involved trimming adapter sequences and removing reads < 30 bp using CutAdapt (Martin, 2011). Overlapping paired-end reads were merged using FLASH (Magoč & Salzberg, 2011). Reads were mapped to the reference genome assembly of the giant panda *(Ailuropoda melanoleuca;* Hu et al., 2017), which represents an outgroup to the investigated clade, using bwa (Heng Li & Durbin, 2009) and samtools (Heng Li et al., 2009), with subsequent filtering for map quality (-q 30) and PCR duplicates (rmdup). These data processing steps were carried out within the BEARCAVE v.1.2 data analysis and storage environment (available at: https://github.com/nikolasbasler/BEARCAVE), which provides a convenient resource for data processing and the establishment of a common sequencing data repository. The specific BEARCAVE scripts used were: *“trim_merge_DS_PE_standard.sh"* for trimming and merging paired-end data generated from double stranded libraries; *“trim_merge_SS_PE_CL72.sh”* for trimming and merging paired-end data generated from single stranded libraries; *“trim_SE.sh”* for trimming single-end data; “*map_SE_0.01mismatch.sh*” for mapping ancient data (only merged paired-end reads were mapped for ancient datasets, which are effectively single-end); and “*map_modern_PE_0.01mismatch.sh*” for mapping modern paired-end data. All details of software versions and parameters can be obtained from the BEARCAVE v.1.2 distribution, which can also be used to replicate the described analyses.

### 2.3 Generation of the Consensify sequences

To generate a Consensify sequence for each dataset, bases were counted at each position of the reference genome using the -doCounts function in angsd v.9.2.0 (Korneliussen, Albrechtsen, & Nielsen, 2014), filtered for minimum base quality of 30 (-minQ) and minimum map quality of 30 (- minMapQ). Base counts were not collected for scaffolds < 1 Mb in length (-rf). A custom perl script was then used to perform the Consensify consensus calling described in Section 2.1. This script outputs the sequence in the fasta file format with sequence headers matching those of the reference genome, and calculates the number of successfully called positions. We additionally implemented an optional user-specified maximum read depth filter which can be entered as an integer number. Regions of exceptionally high coverage may represent repetitive elements with accumulations of incorrectly mapped reads which can be excluded using this filter. For the purpose of this study, we first calculated the 95^th^ percentile of coverage using the -doDepth function in angsd v.9.2.0 and implemented the integer number below this value as the maximum allowed depth for consensus calling. The number of Consensify sites successfully called for each dataset is reported in Table 1. The Consensify script is freely available on GitHub (http://github.com/jlapaijmans/Consensify).

### 2.2 Effect of Consensify on phylogenetic and clustering analysis

We compared the performance of Consensify and standard pseudohaploidisation on phylogenetic and population clustering analyses based on genetic distances. Genetic distance matrices were computed by standard pseudohaploidisation in angsd v.9.20, filtered for minimum base quality of 30 (-minQ) and minimum map quality of 30 (-minMapQ), excluding scaffolds < 1MB (-rf), and only considering sites with zero missing data (-minInd N) that were below the 95^th^ percentile of global coverage (- setMaxDepth), which was determined in advance using angsd (-doDepth). Three distance matrices were calculated by standard pseudohaploidisation including: 1.) all sites; 2.) transversions only (- rmTrans); and 3.) transversions only with singleton removal (1/N < -minFreq < 2/N). A distance matrix was then calculated from the Consensify sequences by combining them into a multi-sequence fasta alignment excluding all columns with missing data, using a custom bash script (‘ReDuCToR’, available from GitHub: http://github.com/jlapaijmans/Consensify). The distance matrix was calculated under the JC69 substitution model using the dist.dna function in the R package *ape* (Paradis, Claude, & Strimmer, 2004; R Core Team, 2013), considering all sites (both transitions and transversions). Neighbour-joining trees for the four approaches were then calculated using the nj function in *ape*, and rooted using the Asiatic black bear outgroup. For population clustering analysis, distance matrices were re-calculated excluding the Asiatic black bear and principal coordinates analysis carried out using the pcoa function in *ape*.

### 2.3 Effect of Consensify on admixture tests

We investigated the performance of Consensify for admixture analysis using the D statistic, and compared it with both the D statistic calculated using standard pseudohaploidisation (standard D statistic) and the extended D statistic with error correction applied to the ancient datasets. The significance of the D value was assessed using a 5 Mb weighted block jackknife test with Z-scores > 3 being considered as statistically significant. D statistics were calculated from the Consensify sequences using the published C++ script D_stat.cpp, and the results processed using the python scripts D-stat_parser.py and weighted_block_jackknife.py ((Barlow et al., 2018), available from https://github.com/jacahill/Admixture). Standard D statistics were calculated in angsd v.9.20 (- doAbbababal) excluding transition sites (-rmTrans). Sites were further filtered for minimum base quality of 30 (-minQ) and minimum map quality of 30 (-minMapQ), excluding scaffolds < 1MB (-rf), and only considering sites that were below the 95^th^ percentile of global coverage (-setMaxDepth). The standard D statistic results were processed using the R script jackKnife.R, which is included in the angsd distribution. Extended D statistics were also calculated in angsd (-doAbbaBaba2) using the same filters. Error rates in the ancient datasets were estimated using the high quality modern Asiatic black bear dataset as the error free individual and the giant panda genome sequence as outgroup. A majority rule consensus fasta sequence was generated from the Asiatic black bear bam file with map and base quality filters (30) using angsd (-doFasta 2) prior to error estimation. Error rates were then estimated for each ancient sample relative to this high quality consensus sequence in angsd (- doAncError), considering only scaffolds > 1 Mb with map and base quality (30) filters applied. The error correction was applied to the extended D statistic ABBA and BABA counts using the R script estAvgError.R, which is included in the angsd distribution.

We first compared the performance of the three D statistic methods on simulated ancient DNA data. Among the three sampled modern brown bears, the Slovenian and Italian individuals are more closely related to each other than either is to the Swedish individual ((Slovenia,Italy),Sweden). D statistic analysis finds no evidence that the Slovenian and Italian populations are differentially admixed with the Swedish population (Z < 3), and thus provides a suitable null model. We then modified data from the Italian bear *in silico* to mimic specific properties of ancient DNA using the program TAPAS ((Taron, Lell, Barlow, & Paijmans, 2018), available from https://github.com/mlell/tapas). Reads were first trimmed to either 35 bp or 50 bp in length using skewer ((Jiang, Lei, Ding, & Zhu, 2014)), and TAPAS was used to introduce C→T substitutions around the sequence ends with a proportion of 0.3 at the terminal nucleotides decaying exponentially towards the median nucleotide, and increase the global misincorporation rate (e.g. sequencing error) by 0.1%. The simulated ancient sequences were then mapped using BEARCAVE v.1.2 and substituted for the unmodified Italian bear data to investigate the effect on D statistic analysis. These tests were run using both the polar bear (SRS412584) and the Asiatic black bear as outgroup.

We then assessed the effect of library preparation method on D statistics by using the double- and single-stranded *ingressus* cave bear datasets as P1 and P2, respectively, with all other cave bears as P3 and the Asiatic black bear as outgroup. We additionally tested for admixture among all combinations of cave bear compatible with their species tree, for admixture between all cave bears and the brown bear lineage (represented by the Slovenian individual) subsequent to the divergence of brown bears and polar bears (represented by individual SRS412584), and for differential brown bear admixture among all cave bear pairs. These tests used the Asiatic black bear as outgroup.

## 3. RESULTS

### 3.1 Statistical properties of the Consensify method

Application of the simple model of sequencing error revealed key properties of the Consensify method. For both Consensify and standard pseudohaploidisation, error rates are lower for heterozygous positions than for homozygous positions, but in both cases Consensify gives substantially lower error rates overall (Fig. 1a). For standard pseudohaploidisation error rates scale linearly with global sequencing error, but for Consensify the error rate scales exponentially (Fig. 1a). As a result, although the absolute difference in error rates provided by the two methods increases with global sequencing error (Fig. 1a), the ratio between them reduces (Fig. 1b). For example, under the assumptions of the model, Consensify provides an approximately 130-fold reduction in error rate compared with standard pseudohaploidisation at a global error rate of 1%, and an approximately 27-fold reduction at a global error rate of 5% (Fig. 1b).

**Figure 1.**
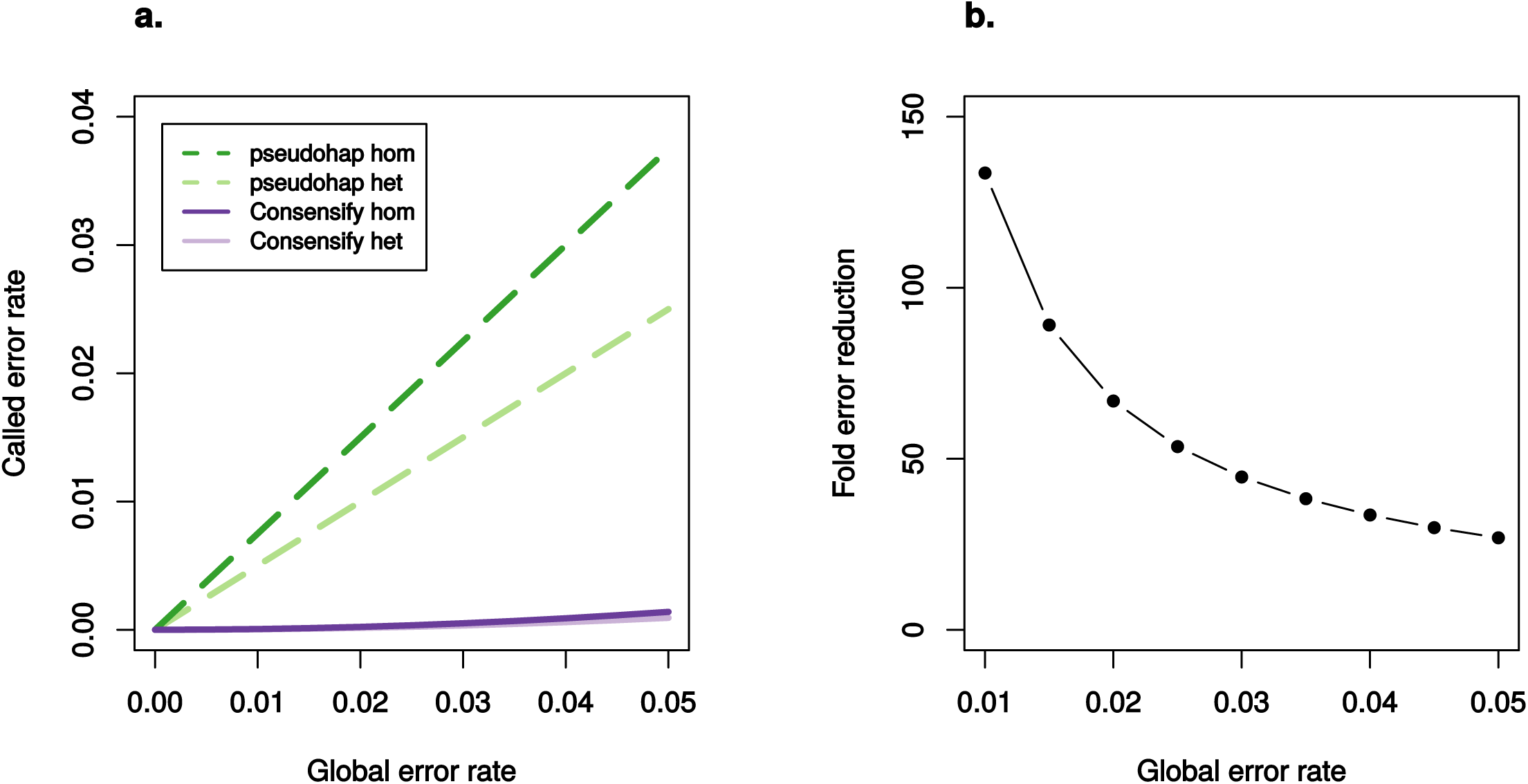
Expected performance of Consensify compared with standard pseudohaploidisation, assuming equal base composition and equal error probabilities across all nucleotides. (a.) shows the expected called error rates (y axis) across a range of global error rates (x axis) for standard pseudohaploidisation (dashed coloured lines) and Consensify (solid coloured lines), for both homozygous sites (dark coloured lines) and heterozygous sites (pale coloured lines). (b.) shows the fold-reduction in error rates achieved by using Consensify compared with standard pseudohaploidisation (y axis), for a range of global error rates (x axis).

### 3.2 Effect of Consensify on phylogenetic and clustering analysis

Distance matrices used for neighbour-joining phylogenetic analysis were calculated by standard pseudohaploidisation from 328,674,048; 244,930,422; and 591,794 filtered variable positions for the all sites, transversions only and transversions only with singleton removal treatments, respectively. The alignment of Consensify sequences included 131,534 filtered variable sites. Phylogenetic analysis recovered the expected topology for all treatments, however differences in branch lengths between treatments were evident (Fig. 2). Using all sites suggested clocklike evolution for the polar-brown bear clade, but cave bear branches are extremely long and variable, consistent with increased and differential rates of error (Fig. 2a). Filtering for transversions only produced a similar pattern but with less extreme branch lengthening for the cave bears (Fig. 2b). Additionally, the double-stranded *ingressus* dataset, which represented the longest terminal cave bear branch when using all sites, is the shortest terminal cave bear branch in the phylogeny calculated from transversions only. Using transversions only with singleton removal produced a phylogeny with more clocklike evolution overall, but with evident branch shortening effects on the more divergent terminal lineages, such as the three brown bear lineages and the *kudarensis* cave bear lineage (Fig. 2c). Analysis of the Consensify sequences produced the phylogeny with the most clocklike evolution, with all tips approximately aligned except for the double-stranded *ingressus* dataset, for which a moderate branch lengthening artefact is evident (Fig. 2d).

**Figure 2.**
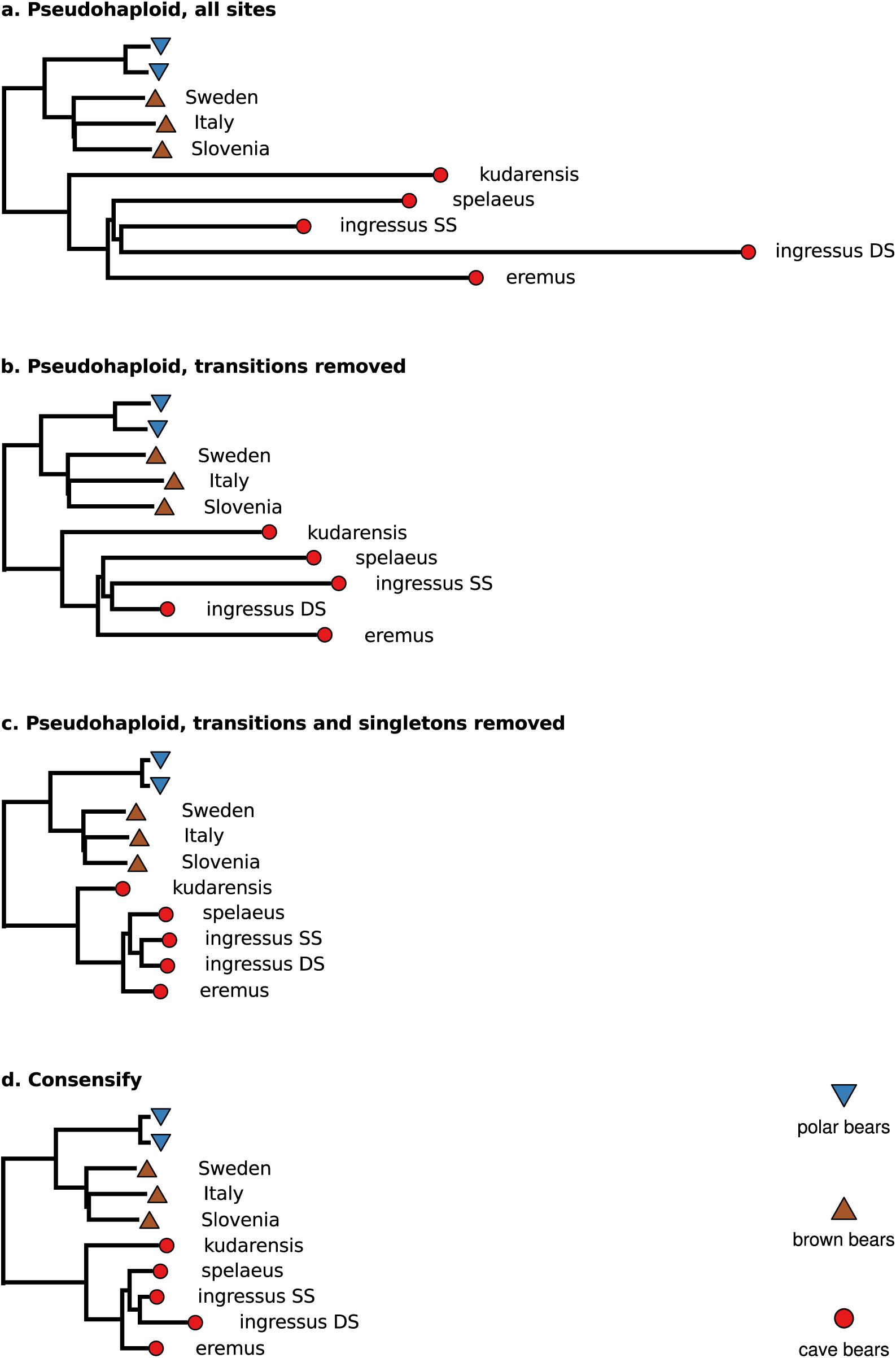
Effect of Consensify on phylogenetic analysis. Panels show neighbour-joining phylogenetic trees calculated by (a.) standard pseudohaploidisation using all sites, (b.) standard pseudohaploidisation with transitions removed, (c.) standard pseudohaploidisation with transitions and singletons removed, and (d.) Consensify. The trees are rooted using the Asiatic black bear outgroup (not shown). Coloured symbols at the terminal tips indicate polar bears (blue triangles), brown bears (brown inverted triangles), and cave bears (red circles). The sampling localities of brown bears and the taxon names of cave bears are indicated. Note that the ingressus cave bear is represented twice, corresponding to datasets generated from sequencing libraries prepared using a single-stranded (SS) and a double-stranded (DS) protocol, respectively. Absolute branch lengths are not comparable among trees because each dataset includes different numbers of sites filtered in different ways. To improve visualisation of relative differences in branch lengths, the trees have been scaled so that the distance between the basal ingroup node and the terminal tips of the polar bear lineage are approximately equal. Polar bears show low genomic diversity (Cahill et al., 2013) and are approaching complete lineage sorting (Barlow et al., 2018), and thus represent the most stable element of the phylogeny with which to anchor the scaling of the trees.

Distance matrices used for population clustering analysis were calculated by standard pseudohaploidisation from 341,311,891; 255,078,638; and 554,431 filtered variable positions for the all sites, transversions only and transversions only with singleton removal treatments, respectively. The alignment of Consensify sequences included 114,891 filtered variable sites. Ordination of individual datasets along the first and second principal coordinates revealed substantial differences between treatments (Fig. 3). Using all sites resulted in separation of the double-stranded *ingressus* dataset from all other individual datasets along the first principal coordinate, and the separation of all other cave bears datasets along the second principal coordinate (Fig. 3a). Polar and brown bear datasets are approximately overlaid, suggesting that the overall pattern is driven by excessive error rates in the cave bear datasets. Filtering for transversions only similarly separated the cave bear datasets along the first and second principal coordinates, with all polar and brown bear datasets approximately overlaid, but the separation of the double-stranded *ingressus* dataset is less extreme (Fig. 3b). Using transversions only with singleton removal produced three clusters corresponding, respectively, to cave bears, brown bears, and polar bears (Fig. 3c). Within the cave bear cluster, the *kudarensis* cave bear is distinct from the other cave bear datasets. Overall, this pattern matches with expectations based on phylogeny (Fig. 2). Analysis of the Consensify sequences produced a similar pattern of three clusters corresponding, respectively, to cave bears, brown bears, and polar bears (Fig. 3d). However, within the cave bear cluster, the double-stranded *ingressus* dataset is distinct from the single-stranded cave bear datasets.

**Figure 3.**
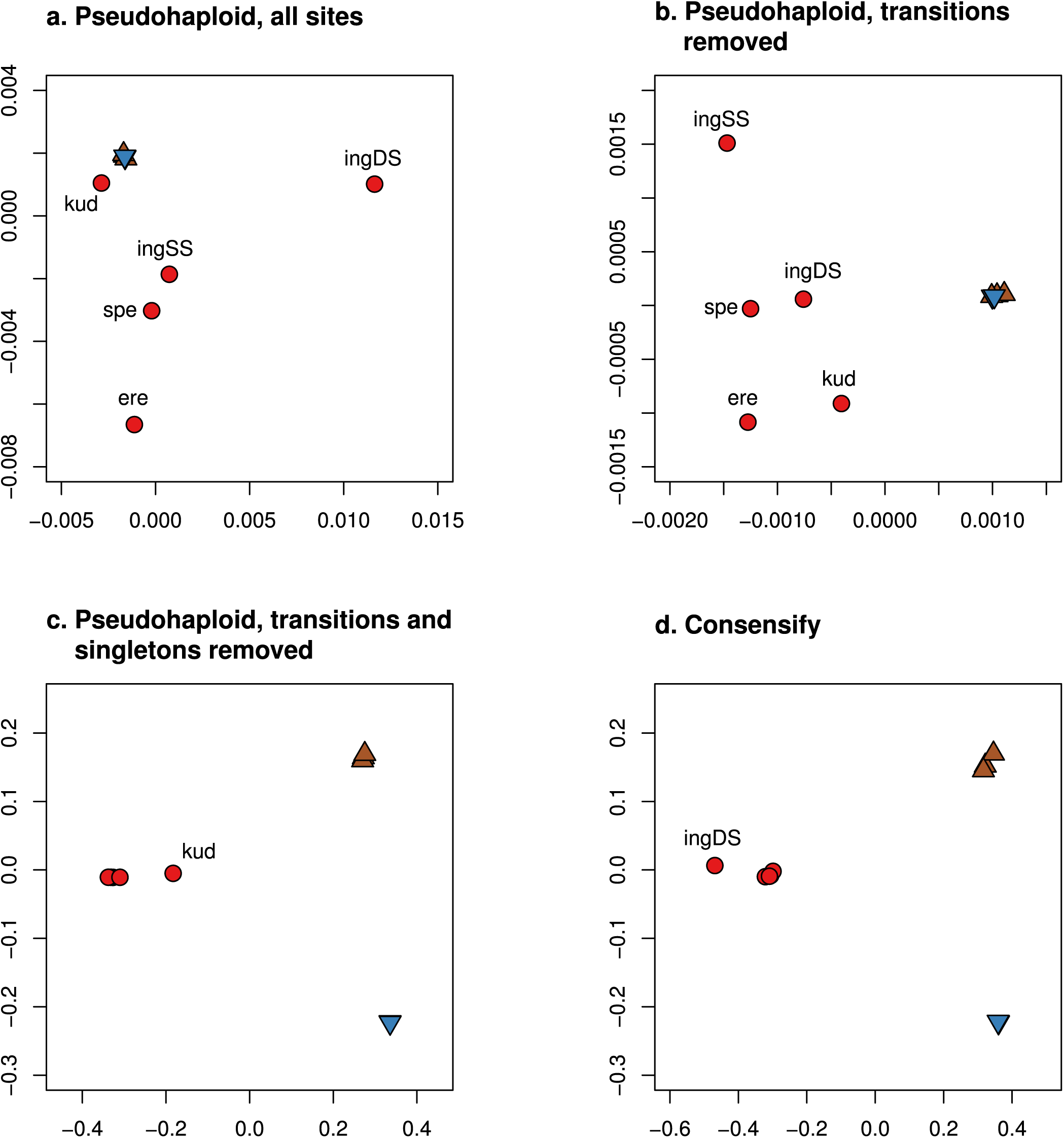
Effect of Consensify on population clustering analysis. Panels show the ordination of individuals along the first (x axes) and second (y axes) coordinates of a principal coordinates analysis based on (a.) standard pseudohaploidisation using all sites, (b.) standard pseudohaploidisation with transitions removed, (c.) standard pseudohaploidisation with transitions and singletons removed, and (d.) Consensify. Coloured symbols are consistent with Figure 1, and where appropriate individual cave bears are indicated by the first three letters of their taxon name.

### 3.3 Effect of Consensify on admixture tests

D statistic tests of admixture among brown bears using the unmodified Italian brown bear data produced non-significant D values across all three D statistic methods, for both polar bear and Asiatic black bear outgroups (Fig. 4). In these comparisons, Consensify recovered a larger number of D statistic informative sites than either the standard D statistic or the extended D statistic, presumably as a result of these datasets having reasonable coverage and because the Consensify analysis makes use of both transitions and transversions, whereas the other methods use only transversions. Substitution of the unmodified Italian bear data with simulated ancient DNA data with 50 bp fragment length, ancient DNA damage and sequencing error also produced non-significant D values across all three D statistic methods for the polar bear outgroup (Fig. 4a), but with the Asiatic black bear outgroup the standard and extended D statistics both produced significant positive D values whereas the Consensify D-value remained non-significant (Fig. 4b). This result does not appear to be driven by a loss of statistical power using the Consensify method, since the number of D statistic informative sites sampled by each method is approximately equal. Analysis of simulated ancient DNA data with 35 bp fragment length using the standard and extended D statistics both produced significant positive D values across all treatments for the polar bear outgroup, but the Consensify D values remained nonsignificant (Fig. 4a). With the Asiatic black bear outgroup, analysis of simulated 35 bp ancient sequences produced significant positive D values for all treatments, but the Consensify D values were closer to zero and with lower Z scores than obtained using either the standard or the extended D statistic. (Fig. 4b).

**Figure 4.**
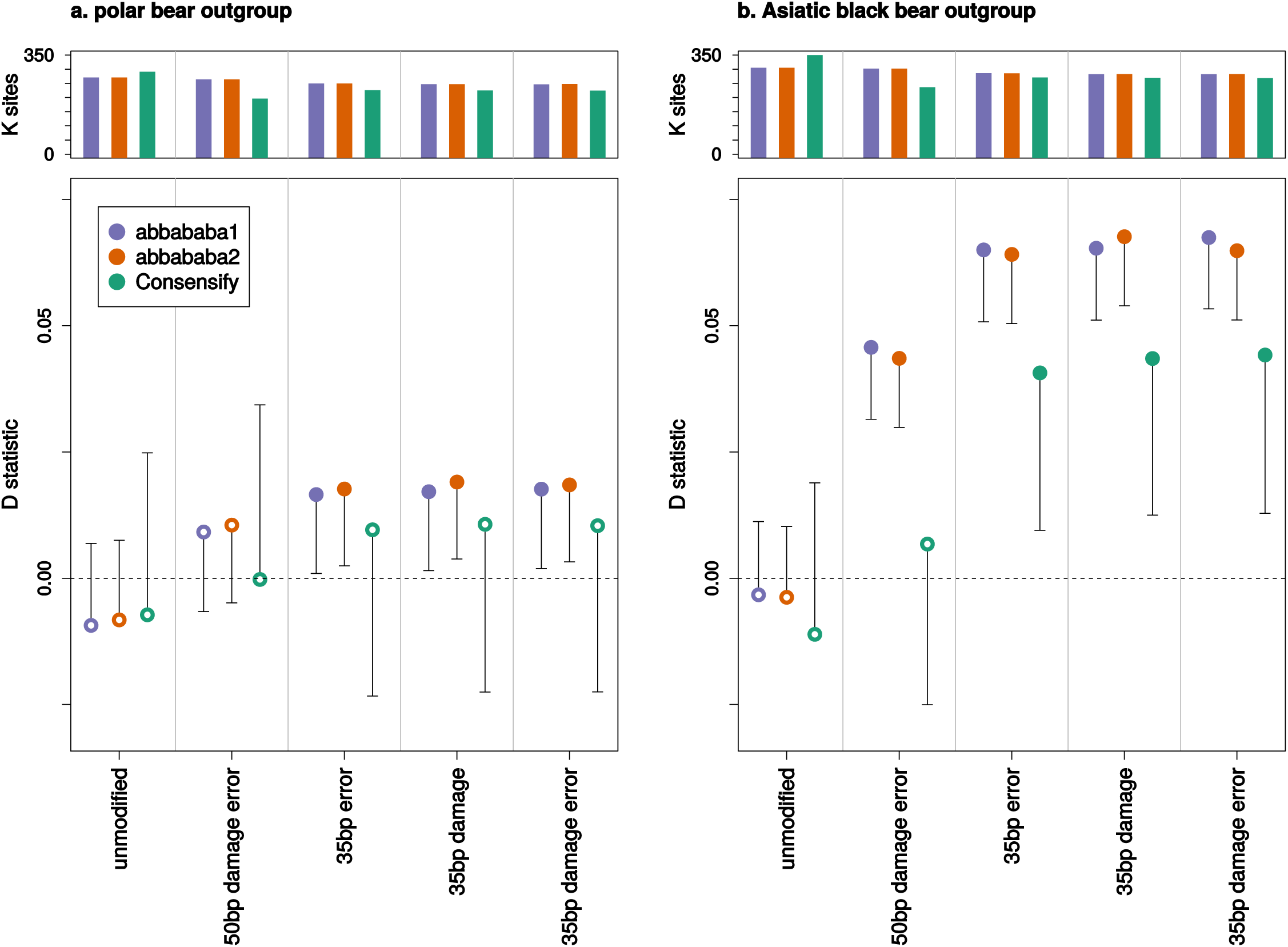
Effect of Consensify on D statistic tests of admixture, evaluated using simulated palaeogenomic data. The tests are based on three brown bears with the relationship: (((P1=Italy,P2=Slovenia),P3=Sweden),P4=outgroup). Each panel displays results calculated using different outgroups: the closely related polar bear (a.) and the more distantly related Asiatic black bear (b.). The upper plot of each panel shows the number of D statistic informative sites (ABBA+BABA, y axes in thousands of sites) counted for each D statistic comparison (separated by grey vertical lines). For each comparison, three results are displayed sequentially from left to right, corresponding to the D statistic (abbababal, purple), the extended D statistic with error correction (abbababa2, orange), and the D statistic calculated using Consensify (green). The lower plots show D values (y axes) as coloured points. Single error bars extending toward zero show the weighted block jackknife standard error multiplied by three, with error bars that bisect y=0 (dashed horizontal line) being non-significant (Z < 3, open points). Significant positive D values are indicated by filled points and may be interpreted as admixture between the Slovenian and Swedish brown bear populations subsequent to the divergence of the Italian and Slovenian brown bear populations. The leftmost comparison in each of the panels corresponds to the original, high quality datasets, and does not provide evidence of admixture in any test. For each adjacent comparison, data from the Italian brown bear has been modified in silico to mimic specific properties of palaeogenomic datasets (x axes): short fragment length (35 or 50 bp), C→T substitutions increasing exponentially towards the terminal fragment ends

Comparisons of the double- and single-stranded *ingressus* cave bear datasets as P1 and P2, respectively, with other single-stranded cave bear datasets as P3, produced significant positive D values using all three methods (Fig. 5a). No method produced obviously lower D values, although standard errors were larger using Consensify and fewer D statistic informative sites were sampled.

**Figure 5.**
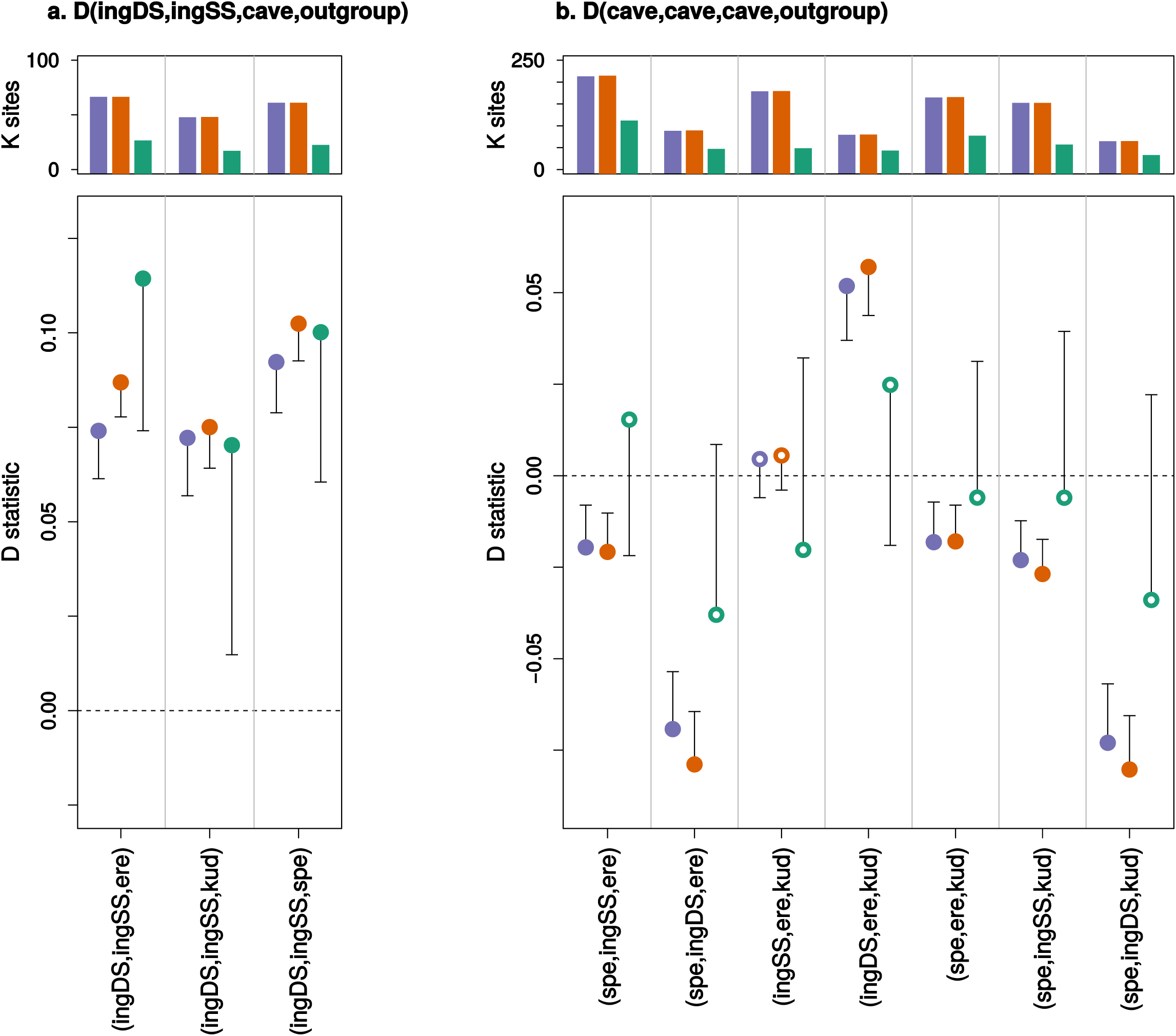
Effect of Consensify on D statistic tests of admixture among cave bear populations and datasets. The plot layout and annotation is consistent with Figure 4. Comparisons are described by x axis labels, with the first three letters of each cave bear taxon indicating their respective positions as (P1,P2,P3). The outgroup (P4) is the Asiatic black bear. The left panel (a.) shows comparisons with datasets generated from the same ingressus cave bear individual as P1 and P2, corresponding, respectively, to datasets generated using either the single-stranded (SS) or the double-stranded (DS) library protocol. The right panel shows all comparisons compatible with the cave bear phylogeny (see Figs. 1 and 3): (((ingressus,spelaeus),eremus),kudarensis). Note that y axes are not consistent between panels.

Admixture tests among all combinations of cave bears compatible with their species tree using the standard and extended D statistics produced significant non-zero D values for all but one comparison (Fig. 5b). This single non-significant value tested for differential admixture with the *kudarensis* lineage among *eremus* and the single-stranded *ingressus* dataset. It is notable, however, that substitution with the double-stranded *ingressus* dataset in this test produced significant positive D values using both the standard and extended D statistics, and a general effect of increased D values associated with the double- vs. single-stranded *ingressus* datasets was apparent across all tests. D values calculated using Consensify were non-significant for all comparisons, and closer to zero for all comparisons where the standard and extended D values were significant.

Compatible with previous studies (Barlow et al., 2018), tests of admixture between cave bears and brown bears subsequent to the divergence of polar bears and brown bears were significant using all three methods (Fig. 6a). Tests for differential brown bear admixture among all cave bear pairs in general supported a geneflow event subsequent to the divergence of *kudarensis* and the European cave bear clade *(ingressus, spelaeus, eremus)*, but with two of these comparisons being non-significant using Consensify and one using the standard D statistic (Fig. 6b). Both standard and extended D statistics supported an additional geneflow event into the *ingressus* lineage, but only for tests involving the single-stranded *ingressus* dataset. All tests among European cave bears were non-significant using Consensify.

**Figure 6.**
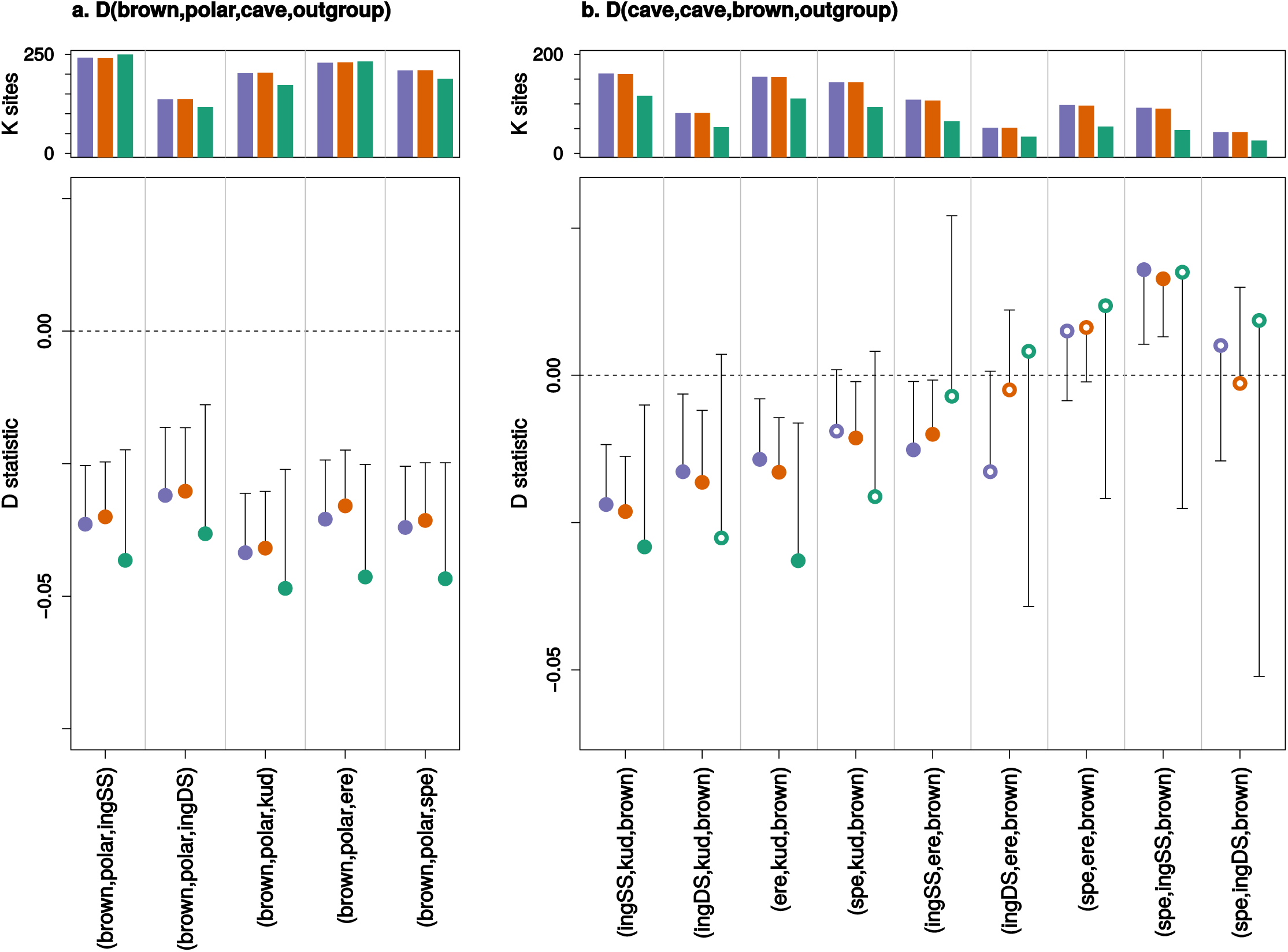
Effect of Consensify on D statistic tests of admixture among cave bears and brown bears subsequent to the divergence of polar bears and brown bears (a.), and subsequent to the divergence of the sampled cave bear populations (b.). The plot layout and annotation is consistent with Figures 4 and 5. The polar bear and brown bear lineages are each represented by a single individual (SRS412584 and 191Y Slovenia, respectively). Comparisons are described by x axis labels, with either the first three letters of each cave bear taxon, or “polar” for the polar bear and “brown” for the brown bear, indicating their respective positions as (P1,P2,P3). The outgroup (P4) is the Asiatic black bear. Note that y axes are not consistent between panels.

## 4. DISCUSSION

High error rates in palaeogenomic datasets are intrinsic since they appear as a direct result of the physical properties of the ancient DNA molecules. Methods of reducing these errors are therefore likely to remain a key aspect of ancient DNA research. Consensify achieves this by normalising coverage bias while leveraging the improved accuracy of calling a consensus from multiple reads. For the datasets analysed here, we have shown that Consensify produces fewer analytical artefacts across a range of methods than observed with other frequently used approaches.

Compared to standard pseudohaploidisation, Consensify produced phylogenetic branch lengths which fitted closer with molecular clock expectations (Fig. 2). Although the removal of transitions and singletons from standard pseudohaploid sequences also produced reasonable trees, the branch reduction artefacts associated with undersampled and divergent lineages may be undesirable. This effect could be mitigated by careful sampling, but this may not be possible in all cases and is a difficult solution to implement *a priori*. Consensify does not suffer such artefacts and may therefore be better suited for analyses with unbalanced or unknown sampling of clades.

One aspect of the test datasets that Consensify failed to fully mitigate are differential errors among single- and double-stranded datasets. Artificial divergence was obvious both with phylogenetic and with population clustering analyses, above that occurring with the transition and singleton removal treatment (Figs 2 & 3). It is feasible that removing transitions from the Consensify sequences may improve this result, but such an approach would dramatically reduce the number of recovered sites when sequencing coverage is low. Currently, all individual cave bears with sequenced genomes are represented by datasets generated using single-stranded libraries. Thus, it is possible to analyse their evolutionary relationships using Consensify from highly consistent datasets generated using identical methods (Fig. 7). The resulting phylogenetic tree shows very clocklike evolution and no branch shortening artefacts as found with singleton removal (Fig. 7a). Population clustering returns three distinct groups corresponding, respectively, to the sampled major bear clades (Fig. 7b). Although consistency of laboratory protocols thus provides an effective solution, implementing this solution retrospectively for published ancient datasets generated using varying library preparation as well as DNA extraction methodologies would represent a substantial challenge.

**Figure 7.**
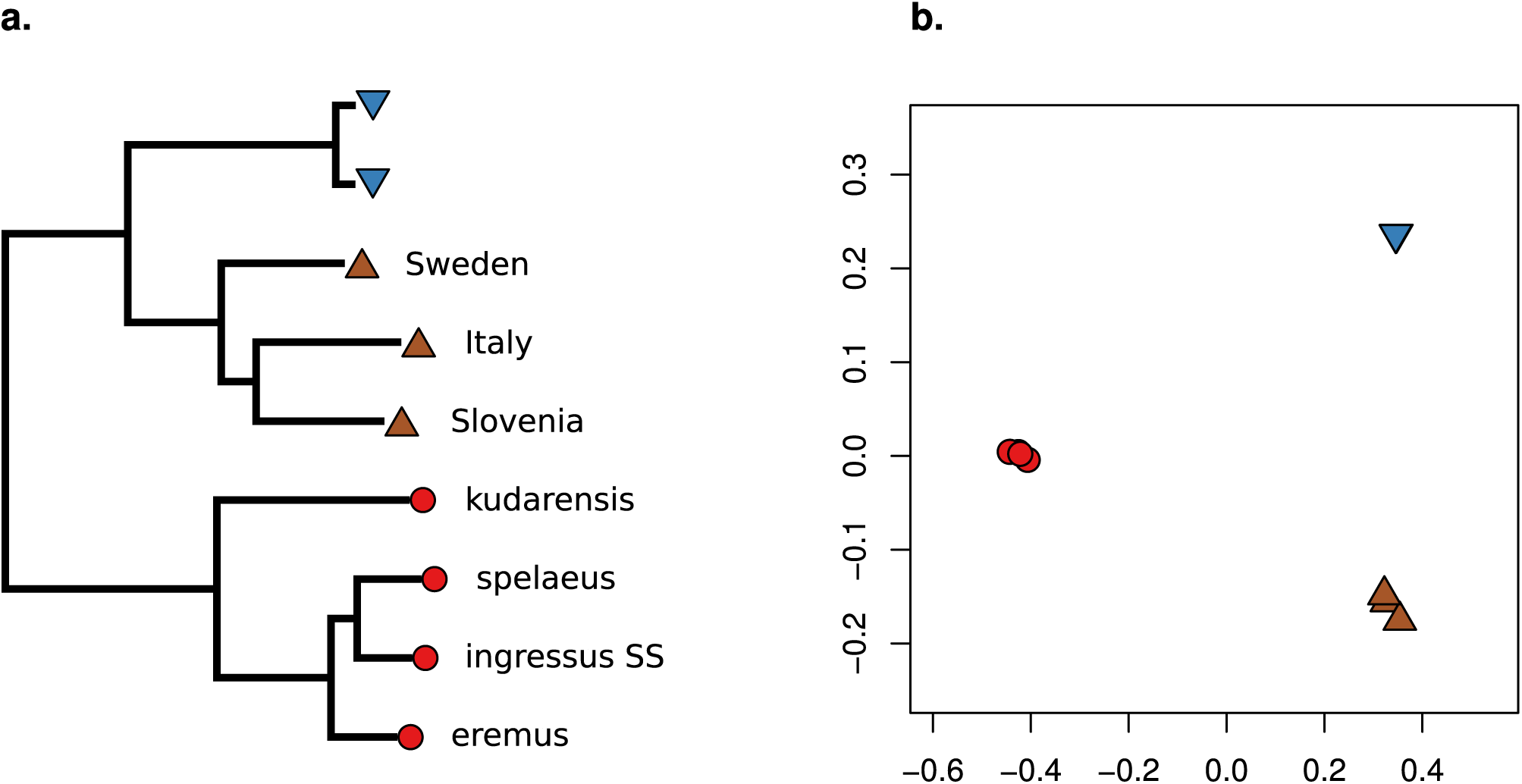
Evolutionary relationships among bears estimated using Consensify. For these analyses, the ingressus cave bear dataset generated using the double-stranded library protocol (ingressus DS) has been excluded to achieve consistency of methods across all cave bears, (a.) Maximum-likelihood tree assuming a phylogenetic model of evolution and a GTR+GAMMA model of nucleotide substitution, rooted using an Asiatic black bear outgroup (not shown). Coloured symbols and tip labels are consistent with Figure 1. (b.) Ordination of the same individuals along the first (x axis) and second (y axis) coordinates of a principal coordinates analysis.

Our results indicate a profound effect of differential error rates on D statistics. Based on the analysis of simulated ancient data, the fragment length seems to be the dominant driver of false positives, having a greater effect on D values than the tested levels of cytosine deamination and global sequencing error (Fig. 4). This would suggest that a large proportion of errors in ancient DNA datasets results from short fragments being incorrectly mapped, although further investigation would be required before this hypothesis can be strongly supported. Nonetheless, across all simulated ancient DNA treatments, Consensify was more resistant to false positives than both standard and extended D statistics. One factor in the generally more conservative results using Consensify is an increase in standard error values compared with standard and extended D statistics. Although Consensify often sampled fewer D statistic informative sites, absolute numbers were generally in the tens to hundreds of thousands. Thus, non-significant results would not appear to result from insufficient statistical power. This is further supported by the fact that the Consensify D values are always closer to zero in false positive tests using simulated ancient data than the standard and extended D values (Fig. 4). We suspect that the increased standard errors may instead reflect the patchy mapping of reads to the divergent panda reference genome, which will be exacerbated at low coverage when only regions with a read depth above two or three are selected using Consensify. This would lead to greater variance when any single 5 Mb clock is removed for weight block jackknife analysis. If this is the case, mapping to a closely related reference (e.g. ancient human data to the human genome assembly) should not produce such large standard errors, but this is currently untested.

Further support for the utility of Consensify is provided by D statistic tests of admixture among cave bears. Of all tests performed, these are most likely to be affected by differential errors as all ingroup individuals are ancient. In line with this, the standard and extended D statistics returned significant values for all but one comparison among cave bears (Fig. 5b). If correct, many of these inferred geneflow events are difficult to explain. For example, cave bears from the Caucasus Mountains *(kudarensis)* would have to admix with those in the west of Europe *(spelaeus)* to a greater extent than the geographically more proximate cave bear populations in eastern and central Europe *(ingressus)*. The inferred occurrence of admixture between *kudarensis* and *ingressus* also changes depending on whether double- or single-stranded datasets are used. Using Consensify, no such complex interpretations are required as no significant evidence of admixture is found among *kudarensis* and any one of the sampled European cave bear lineages, or among *eremus* and either *spelaeus* or *ingressus*, which is compatible with evidence from mitochondrial DNA (Hofreiter et al., 2004; Stiller et al., 2014).

It is surprising that in false positive tests on simulated ancient data, the performance of the extended D statistic was not especially different to the standard D statistic (Fig. 4). This was unexpected, since the extended D statistic in theory provides a true error correction. Consensify, by contrast, only reduces the absolute error rate, meaning that the relative difference in error rates among samples likely remains similar. Consensify therefore relies on reducing differences in ABBA and BABA counts occurring due to errors substantially below that occurring due to population processes. This effect is evident from comparisons with the double- and single-stranded *ingressus* cave bear datasets as P1 and P2, where D values were similar and significant across all methods (Fig. 5a). Since differences in ABBA and BABA counts in these tests are solely driven by differential errors resulting from different methods of library preparation, their ratio (and the resulting D value) remains largely unchanged using Consensify. When applied in tests among different cave bears, however, Consensify seems to better mitigate the effect of mixed methods of library preparation, since the inferred patterns of admixture were unchanged when the double- and single-stranded *ingressus* datasets were substituted (Fig. 5b). Overall, our results therefore suggest that, at least for the datasets included in this study, Consensify provides lower false positive rates and generally more conservative estimates of admixture than the extended D statistic.

A limitation of Consensify is that the amount of sequencing required to achieve a certain number of pseudohaploid sites will be higher than when using standard pseudohaploidisation. Thus at extremely low coverage Consensify will not be applicable because so few sites are covered by two or three reads. This problem is exacerbated when more than one dataset has low coverage, since the probability that any one site has sufficient coverage across all datasets is even smaller. Consensify does mitigate these issues to some extent by making use of transitions as well as transversions, and at higher levels of coverage this can even lead to an increase in informative sites compared with standard pseudohaploidisation (Fig. 4a). Recent discoveries such as the mammalian petrous bone as a source of high purity ancient DNA (Pinhasi et al., 2015), and improved knowledge of the distribution of contaminant DNA across different bone structures (Alberti et al., 2018; Damgaard et al., 2015) mean that achieving levels of genome coverage suitable for Consensify is increasingly possible. For example, each single-stranded ancient dataset analysed here was generated using relatively modest sequencing effort, being approximate to a single lane of sequencing on a current Illumina HiSeq platform (Barlow et al., 2018). For these samples, this produced 3.7 - 6.9 Gb of mapped data resulting in 0.9 - 1.3 Gb of Consensify sequence, providing over 100,000 variable sites after strict filtering for phylogenetic and population clustering analysis, and generally tens of thousands of D statistic informative sites. Until the cost of sequencing reduces to a point where all ancient samples with sufficient surviving DNA can be sequenced to very high coverage, Consensify should represent a useful tool for the analysis of palaeogenomes.

## ACKNOWLEDGMENTS

We would like to thank Nikolas Basler for management of the BEARCAVE environment. We would further like to thank J.J. Paijmans for coding support for the ReDuCToR script. We also thank Ioana N. Meleg for beta testing of the scripts. J.L.A.P. was funded by the University of Leicester (M38DF64). This work was funded by European Research Council (ERC) consolidator grant ‘gene flow’ 310763 to M.H.

## DATA ACCESSIBILITY STATEMENT

The scripts for running the method presented in this manuscript are available on GitHub (http://github.com/jlapaijmans/Consensify). GenBank SRA accession numbers for the data generated for the *ingressus* cave bear (GS136_ss) will be made available upon acceptance.

## AUTHOR CONTRIBUTIONS

AB and JLAP invented and developed the Consensify method; SH wrote the Consensify perl script; JLAP wrote the ReDuCToR script for generating Consensify alignments; JG performed lab work; AB and JLAP analysed the data; AB, JLAP and MH interpreted the results; AB and JLAP prepared documentation and assembled the Consensify v.0.1 distribution; AB wrote the manuscript with input from all coauthors.

